# Epidemiological, Serological, and Virological Analysis of an Outbreak of Elephant Hemorrhagic Disease in Switzerland

**DOI:** 10.1101/2024.03.14.585086

**Authors:** Mathias Ackermann, Jakub Kubacki, Sarah Heaggans-Ebbeson, Gary S. Hayward, Julia Lechmann

## Abstract

Elephant hemorrhagic disease (EHD), caused by several Elephant endotheliotropic herpesviruses (EEHV), represents a frequently lethal syndrome, affecting both captive and free-living elephants. In summer 2022, three young Asian elephants (*Elephas maximus*) succumbed to EHD in a zoo in Switzerland, despite of considerable preventive efforts and early detection of EEHV1A viremia. In this communication, we describe the extent of preventive measures in terms of prior virus detection, active survey of viremia, and antibody status. In the course of the outbreak, the causative virus was concomitantly analyzed and eventually fully sequenced and compared to other EEHV types and strains.

The conclusions from these analyses may be summarized in three points: (1) A previously undetected EEHV1A strain had remained unrecognized among these elephants. Probably, the new virus re-emerged after almost 40 years of latency from one of the oldest elephants in the zoo. (2) While two of the three affected animals had prior immune responses against EEHV1, their strain-specific immunity proved insufficient to prevent EHD. (3) There is an urgent need to develop efficient antiviral drugs and protective vaccines. In particular, ways need to be found to circumvent the present unavailability of appropriate cell cultures and animal models.

## Introduction

Elephant hemorrhagic disease (EHD) can be caused by several Elephant endotheliotropic herpesviruses (EEHV) and represents a frequently lethal syndrome, affecting both captive and free-living elephants (1-4). Pairs of distinct but genetically related types of EEHV exist in Asian (*Elephas maximus*) and African elephants (*Loxodonta africana*), respectively (5, 6). EEHV type 1 (EEHV1), which has been most frequently associated with the occurrence of EHD in captive Asian elephants, can be further divided into the subtypes A (EEHV1A) and B (EEHV1B) and into so-called ORF-Q clades (4, 6, 7). Clades A, B, C, and D have been firmly established, both on the genetic as well as on the serologic level (7). However, the existence of at least three further clades (E1, E2, and F subtypes) has been documented by GenBank entries (E1: QBP78682; E2: QOE74866.1 and QOE74629; F: QBP78717).

The genetic counterpart of EEHV1 in African elephants has been named EEHV6 (8). Among others, EEHV6 has been linked with cases of EHD in African elephants at the Schönbrunn zoo near Vienna. The other two pairs of EEHV species are represented by EEHV2 in African elephants and EEHV5 in Asian elephants as well as EEHV3 (in *Loxodonta africana*) and EEHV4 (in *Elephas maximus*)(1, 6, 9).

The study of epidemiology, pathogenesis, treatment, and prevention of EEHV-related diseases has been hindered because, as yet, none of the EEHV types, species, and strains can be propagated in conventional cell cultures. Moreover, animal models for EEHV and EHD do not exist because no susceptible animals, except elephants, have ever been detected. Not surprisingly, it took a very long time until full EEHV genomic sequences became available (10, 11).

As with all herpesviruses, the reservoir of EEHV is in latently infected animals, which may intermittently reactivate and shed virus for transmission (1, 12). All known EEHV strains replicate primarily in the respiratory tract but, for poorly understood reasons, can cause viremia, which is the major precondition for developing EHD (13). Another critical parameter for developing EHD is associated to the level of specific anti-EEHV antibodies in the serum. Very young animals are thought to be protected by maternal antibodies under whose cover subclinical infections may take place to induce active immunological protection against the disease in adult animals (7, 14, 15). Accordingly, the highest risk for young animals to succumb to EHD is in the age of between two and eight years and in association with waning of maternal antibodies.

Viral genomic sequences have been very useful to determine transmission chains among elephants, which eventually lead to interconnected cases of EHD. For example, the quite variable E54/EE51 locus (encoding vOX2-1) was shown to be identical between three cases of EHD in Berlin, which occurred 11 years apart: EP14 in the year 2000, EP23 and EP25 in 2011. However, the knowledge of the present viral sequences are also important for prognosis and treatment: subtype 1A is considered the most virulent EEHV affecting Asian elephants, whereas subtype 1B is known to frequently cause milder courses of EHD (1, 16).

Only recently, the value of serology has also been established as a useful tool in the context of EEHV epidemiology as well as in EHD risk management (2, 7, 14, 15).

Albeit highly diverged, all known species of EEHV encode thymidine kinase (Tk) as well as conserved protein kinase enzymes (CPK), i.e. U48.5/EE7 encoding ETk and U69 encoding ECPK, respectively. It has been speculated that these viral enzymes may be capable of phosphorylating nucleoside analogues such as ganciclovir (GCV), aciclovir (ACV), and penciclovir (PCV) or famciclovir (FCV, the PCV-derivate for oral application), which is an essential prerequisite for their potential antiviral activity against EEHV (10, 11, 17-19).

Indeed, nucleoside analogues such as ACV, FCV, and GCV have been used to treat elephants with clinical and virological EHD but the clinical efficacy of these drugs remains at least debatable (1, 20-22), although intravenous treatment with GCV coincided at least in one case with decreasing viremia (23). However, in the absence of animal models and suitable cell cultures, the true value of treatment against EEHV-infections with nucleoside analogues remains unknown (1).

Despite all these uncertainties, the desire to prevent lethal cases of EHD among young elephants in captivity increased with the growing knowledge about its causative agents and their biological properties. Accordingly, many zoos, including ours, initiated surveillance programs to identify prevalent EEHV types and strains and to monitor their animals at risk for early signs of viremia (13, 24). It was expected that typing of the prevalent EEHV strains may help to focus on particularly dangerous isolates, while screening for viremia may allow early onset of antiviral treatment in cases of emergency (16, 20, 23, 25).

Prior to 2022, three cases of EHD had been recorded among Asian elephants in Switzerland, all of them attributable to EEHV1A, the first in 1988 (partial sequences deposited in Genbank as **EP04**, Lohimi: KT705308.1), the second in 1999 (partial sequences in GenBank as **EP07**, Xian: MH287538.1) and the third in 2003 (EP16, Aishu; no sequence entries in GenBank but unpublished U48.5/EE7 sequences, encoding for ETk, classified the isolate as subtype 1A) (24, 26).

In 2014, after many years of successful breeding and, thus, without the need to admitting elephants from outside, a young bull (Thai, see Materials and Methods) was integrated among the elephants in the Zurich zoo, with the aim to replace the aging prior breeding bull (Maxi) and to avoid inbreeding. As a calf had been born in the same year (Omysha), it seemed appropriate to establishing a preventive program, which included risk assessment and active surveillance. Initially, EEHV-shedding was assessed, which resulted in the detection of two additional circulating viruses among these elephants, namely an EEHV1B isolate from Omysha as well as an EEHV4 isolate from Thai (24). Accordingly, the young animals were screened on a weekly basis for early detection of viremia by EEHV1 and EEHV4, respectively (25). Serology was established “*en route*” as it became available from literature (2, 7, 14). Moreover, stocks of antivirals, including GCV and FCV were stored for cases of emergency.

In spite of all these precautions, disaster struck in summer 2022, when three of these young elephants died due to EHD caused by a previously undetected EEHV1A strain.

In this communication, we describe and evaluate our preventive measures and come close to identify the source of the outbreak by serology. With only three animals, our data cannot claim statistical significance but in the absence of suitable cell cultures and animal models, they still help us to improve our understanding and strategies against this dreadful disease.

## Materials and Methods

### Ethical statement

Viremia samples, tissue samples and sera from elephants were analyzed by order of the contributing zoos. Ethical approval was not sought as the present study only made use of samples that were taken for specific diagnostic purposes, the results of which are described in this manuscript. The samples were taken under routine veterinary care by veterinary staff. The contributing zoos as well as their scientific advisors consider it important to sharing critical medical, husbandry, training, and scientific information with colleagues across the globe.

### Elephants

At the time of the outbreak, the herd of Asian elephants (*Elephas maximus*) consisted of eight animals, divided into two family lines and one bull (Thai), the most recent admission to the herd, who had been introduced from outside in 2014 in order to replace the previous breeding bull (Maxi) (Fig 1). The first family line, whose founder Ceyla had been born in 1975 in Sri Lanka and transferred to the zoo in 1976, consisted of three individuals, including the founder’s daughter Farha and her daughter (Ruwani), born to Farha and Thai. As two of Ceyla’s sons (Xian, EP07 and Aishu, EP16) had previously succumbed to EHD, these two individuals were also integrated into Fig 1.

**Fig 1.**
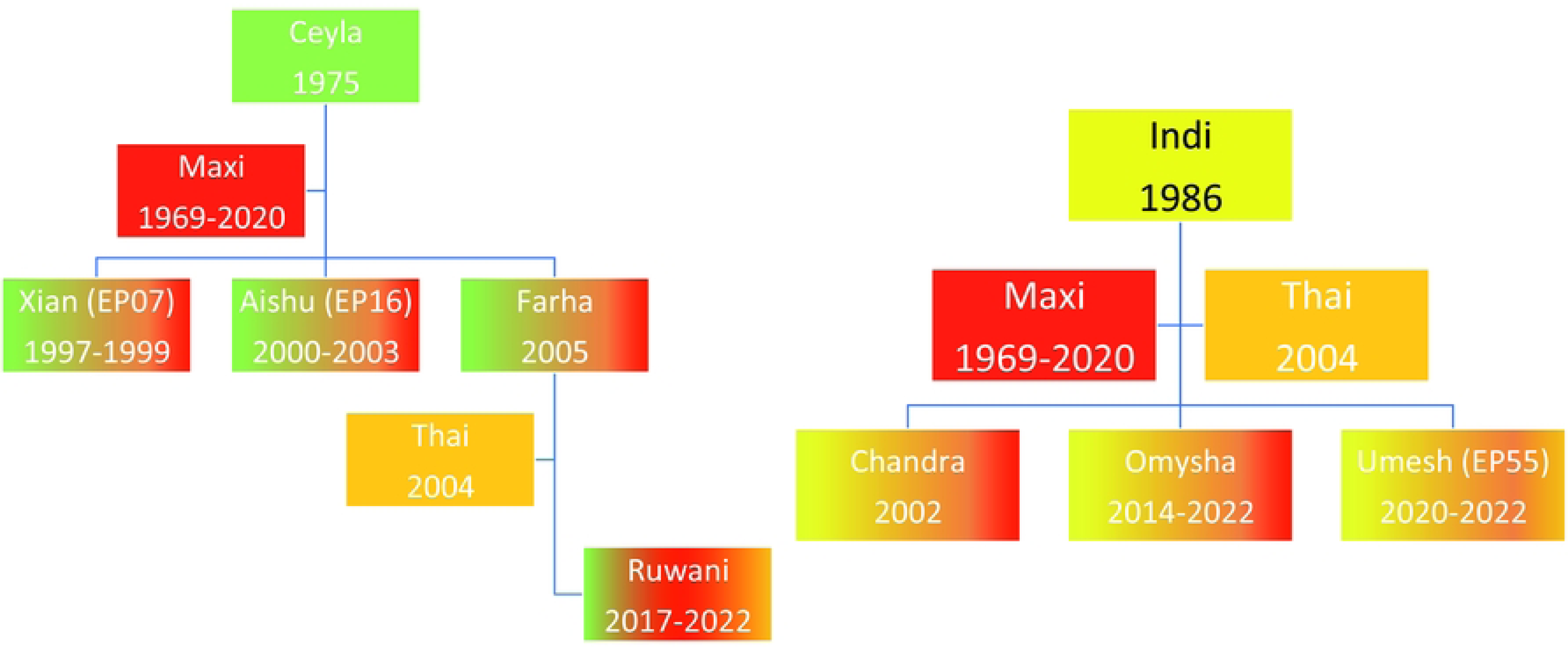
Family trees of the elephants relevant for the present study. Green and yellow: founder females with designation and year of birth. Red and orange: breeding bulls. Multicolored: color coded offspring with designation, year of birth, and year of death (if applicable). Two prior cases of EHD are also integrated.

The second family line, whose founder Indi had been born in 1988 in Myanmar and transferred to the zoo in 1999, consisted of four individuals, including the founder’s two daughters from Maxi (Chandra and Omysha) and her recent son (Umesh) from Thai. Thus, all individuals except of the two founders and the bulls had been born in the Zurich zoo.

### Samples

#### Viremia samples

For routine viremia screens, one blood droplet (approx. 20 µL) was drawn from the elephant’s feet during pedicure by using a small capillary, which then was placed into an Eppendorf tube containing 80 µL nuclease-free water and thoroughly mixed. Historic EDTA blood samples from EHD cases in 2018 (H1 Anjuli and H2 Kanja) were kindly provided by the Hagenbeck zoo in Hamburg, Germany. Cellular fractions were taken as source of DNA extraction for PCR, whereas plasma fractions were used in serology.

#### Serum samples

Sera from Swiss elephants are routinely collected and archived for health monitoring purposes. Samples from these archives were provided for serological analyses by order of the Basel zoo and in the Zurich zoo, respectively. Samples from the Zurich zoo were collected throughout March 2022. Four sera (A1, A2, A3, A4) from African elephants in Basel zoo were collected throughout January 2022.

#### Tissue samples

After necropsy of each of the three elephants, 15 mg from each of five tissues (heart, spleen, liver, kidney, lung; kindly provided by Dr. Monika Hilbe, Institute of Veterinary Pathology, University of Zurich) were combined from each individual and minced in a combination of 300 µL of body fluid and 300 µL blood using a TissueLysser II (Qiagen). In addition, a few slices of historic heart and liver tissues from Xian, son to Maxi and Ceyla, who had been lost to EHD in 1999 (partially sequenced genome in GenBank as EP07), were kindly made available for the present study by Prof. Franco Guscetti, Institute of Veterinary Pathology, University of Zurich.

### Real-time PCR detection of EEHV1, EEHV4, and ETNF

Per well, 5 µL aliquots from diluted blood droplet samples were used as template for detection of EEHV1, EEHV4 and elephant TNF (ETNF), respectively, by real-time PCR as described previously (13, 24). Briefly, each 20 µL reaction volume contained 10 µL PrimeTime Gene Expression Master Mix (Integrated DNA Technologies), 0.9 µM of each primer and 200 nM probe, and 5 µL template or nuclease-free water as negative control. Samples were tested in duplicate on a QuantStudio 3 Real-Time PCR System (Applied Biosystems) using the following cycling conditions: 2 min at 50°C, 10 min at 95°C, 40 cycles of 15 sec at 95°C and 1 min at 60°C.

A test was considered valid if ETNF was detected at a Ct value of 39 or lower and if the negative controls for each assay did not yield a Ct value. To compensate for sample variation in volume and elephant cells, the Ct value of ETNF was used as an approximate measure for the number of elephant cells within the sample and, in positive cases, the Ct value of the ETNF signal was subtracted from the Ct value for EEHV to get a ΔCt value (ΔCt = Ct_EEHV_- Ct_ETNF_). Assuming similar test efficiencies, a ΔCt value of zero signified approximately equal amounts of ETNF and EEHV templates, whereas a positive ΔCt value indicated less than one EEHV genome per cell. Accordingly, the ΔCt values grew on the negative side with increasing numbers of EEHV genomes per cell.

### Conventional PCRs

Two loci on the EEHV1 genome (E36/U79 and E54/EE51 vOX2-1) were targeted by conventional PCR to characterize the current viral genotype very early during the outbreak (Table 1). The PCR conditions were as described previously (24). Briefly, the PCRs were run in a volume of 20 µL, comprising 5 µL template, 10 nM dNTP, 5x Phusion GC buffer, 2 U of Phusion High-Fidelity DNA Polymerase (Thermo Scientific) and each primer at a concentration of 0.5 µM (Table 1). After an initial denaturation for 2 min at 98°C, the samples were cycled 39 times at 98°C for 10 sec, 58.4°C for 20 sec, and 72°C for 7 sec. The runs were completed with 10 min at 72°C before cooling to 10°C. The PCR products were evaluated for their size by agarose gel electrophoresis before being extracted from the gel for cycle sequencing by a commercial company (Microsynth, Balgach, Switzerland).

**Table 1.**
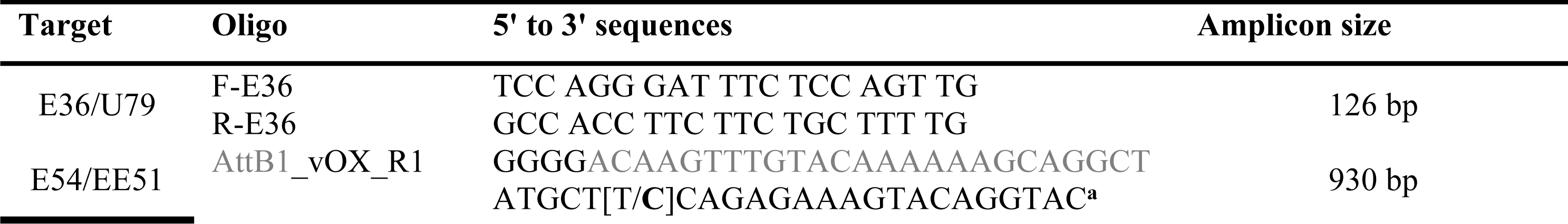
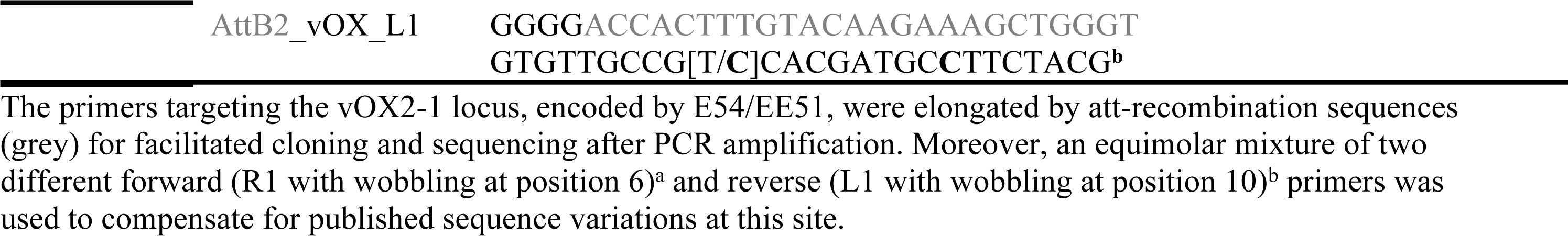
Oligonucleotides used for conventional PCR.

### LIPS assays

#### LIPS antigens

Antigen for LIPS serology was harvested from transfected COS7 cells, using synthetic and codon-optimized DNA provided by GenScript Europe (The Netherlands). For this purpose, the published amino acid sequences of E39 (clades A (Kimba), B (Emelia), C (Raman), and D (Daizy)) as well as of the gB glycoprotein (EEHV1A, Kimba) were reverse translated, codon-optimized for expression in human cells, and cloned under the control of the cmvIE promoter into the pcDNA3.1(+) vector. Moreover, each construct was fused with the following sequence of C-terminal tags: V5-epitope, nanoLuciferase (nLuc), and 7-his. Except for clade D-E39, each construct possessed its native signal peptide, which was included into the gene synthesis design. As clade D-E39 was not predicted to possess a functional signal sequence but two short transmembrane regions close to the N-terminus, the molecule was shortened to start only at its original amino acid No. 60, which was preceded by the clade A signal peptide. Expression of the desired antigens following transfection was verified by measuring nLuc activity (relative light units, RLU) in the cell culture supernatants as well as by detecting the corresponding proteins in cell lysates after standard Western immunoblotting using a commercial anti-V5 monoclonal antibody (SV5-Pk1 from Invitrogen).

#### LIPS assay

The protocol for the LIPS assay was adapted from a previously published overview article (27). A commercial kit (Immunoprecipitation Kit, Abcam), which provided buffers for protein extraction and solubilization as well as Protein A/G sepharose beads and washing buffers, was used according to the supplier’s instructions. Briefly, the secreted antigens were harvested from the cell culture supernatants and mixed with non-denaturing solubilization buffer (NDSB) and protease inhibitors before non-soluble proteins were removed by centrifugation. Prior to the assay, the nLuc activity of each antigen was adjusted to comprise 10^7^ RLU per second (RLU/s) in a volume of 50 µL (NanoGlow, Promega, Madison, Wisconsin, USA).

The sera were prediluted with phosphate buffered saline (PBS) to 1:100 in a polystyrene microtiter plate (Nunc F96 MicroWell flat bottom) and mixed well by shaking. Prediluted serum stocks were kept in these plates at -20°C for a maximum of three weeks. A second polystyrene plate was used for the actual assay. First, 40 µL of NDSB were pipetted into each well before 10 µL of prediluted sera and 50 µL of freshly diluted nLuc-antigen were added. The mixture was incubated with shaking (300 rpm) for one hour at ambient temperature. Meanwhile, a slurry comprising 30% sepharose A/G beads was prepared in PBS and 5 µL were added to each serum sample to incubate for one more hour with shaking. Eventually, all samples were transferred to a filter plate (Merck MultiScreen) and washed on a vacuum manifold. Each well was washed 8 times with 100 μL of wash buffer, followed by two times washing with PBS. Following the last wash, the vacuum was turned off. The filter plate was removed and blotted dry using a stack of filter paper making sure to remove moisture on the top and bottom of the plate. Eventually, 50 µL of NanoGlow substrate (Promega) was added to each well before measuring in a luminometer (MicroLumat Plus, LB 96 V, up-version 2.0; software WinGlow v. 1.25.000003, Berthold Technologies, Bad Wilbad, DE). Two horse sera (P1 and P2), both known to have antibodies against equine herpesvirus 1 (EHV-1) were used as controls to determine the negative cut-off values of each test.

### Sample Preparation and DNA Sequencing

Eventually, DNA was extracted from necropsy samples for whole genome sequencing. A previously established viral particle enrichment protocol was used to increase the recovery of viral genetic material (28). Then, enrichment of viral nucleic acids was followed by reverse transcription for any RNA viruses (to exclude other causes of death) and sequence-independent single primer amplification for DNA. Sheared to 500 bp length using the E220 Focused-ultrasonicator (Covaris, Woburn, MA, USA) and prepared with the NEBNext Ultra II DNA Library Prep Kit for Illumina (New England Biolabs, Ipswich, MA, USA) according to the manual. Sequencing libraries were sequenced at the Functional Genomics Center Zurich (FGCZ) in a paired-end 2 × 150 bp, SP flow cell sequencing run using the NovaSeq 6000 (Illumina, San Diego, CA, USA). The PhiX Control v3 Library (Illumina, San Diego, CA, USA) was added as the control.

### Data analysis

Sequenced reads were analyzed using the de novo assembly pipeline described previously (29). Illumina sequencing adapters and low-quality sequencing ends were trimmed using Trimmomatic (v. 0.39) (30). Second, trimmed reads with average quality above 20 and longer than 40 nt were further cleaned up by trimming away the SISPA primers using cutadapt (v. 3.2). The quality-checked reads were then assembled using metaspades (v. 3.12.0) (31). Assembled de novo contigs were compared to the NCBI non-redundant database (nt) using blastn (v. 2.10.1+) and annotated using the best blastn hits. Quality-checked reads were mapped back to the assembled sequences using bwa (v0.7.17) mem (32). In addition, contigs from EEHVs were aligned in a metagenomic pipeline of the SeqMan NGen v.17 (DNAStar, Lasergene, Madison, WI, USA) to all available viral genomes of other EEHVs downloaded from the NCBI database (08.2022) and visualized in the SeqMan Ultra (DNAStar, Lasergene, Madison, WI, USA).

To determine genome similarity values between our assembled strain and other available complete EEHV1A strains, EEHV1B (Emelia), and EEHV6 (NAP35) we performed BLASTN analysis using the VIRIDIC tool (settings: ‘-word_size 7 -reward 2 -penalty -3 - gapopen 5 -gapextend 2’) (Moraru et al., 2020).

### Sequence sources

Publicly available sequences from NCBI were used for comparison of our data. The most relevant cases are listed in Table 2.

**Table 2.** List of EEHV sequences.

| Virus | Case | Individual | Origin | Year | ORF-Q | Source |
| --- | --- | --- | --- | --- | --- | --- |
| EEHV1A | NAP23 | Kimba | USA | 2004 | A | KC618527.1 |
| EEHV1A | NAP72 | Daizy | USA | 2015 | D | KX759150.2 |
| EEHV1A | EP24 | Raman | UK | 2009 | C | KC462165 |
| EEHV1B | EP18 | Emelia | UK | 2006 | B | KC462164 |
| EEHV1A | EP14 | Plai Kiri | Germany | 2014 | C | MH287551.2 |
| EEHV1A | EP07 | Xian | Switzerland | 1997 | D | MH287543.2<br>MH287538.1 |
| EEHV6 | NAP35 | Miss Bets1 | USA | 2009 | 6 | MZ822422 |

## Results

### Weekly screening for EEHV-viremia

Starting from 21 January 2018, capillary blood was taken on a weekly basis and analyzed by TaqMan real-time PCR for the presence of EEHV1 and EEHV4 DNA, respectively. Initially, only animal Omysha was monitored but the younger animals were included once they tolerated the sampling procedure. The results of this monitoring program are summarized in **Table 3**. Although rarely and at low titers, transient and subclinical viremia by EEHV1 was detected in two of the three monitored animals. Notably, EEHV4 DNA was never detected throughout the screening period.

**Table 3.** Results of TaqMan real-time PCR targeting EEHV1, EEHV4 and ETNF.

| Identification | Screening | Detection of EEHV |
| --- | --- | --- |
| **Omysha** | 231 weeks | 4 times EEHV1; $\Delta Ct > 4$ |
| **Ruwani** | 209 weeks | 3 times EEHV1; $\Delta Ct > 4$ |
| **Umesh** | 60 weeks | Never detected |

### EEHV serology using LIPS assay

A LIPS assay using the conserved glycoprotein B of EEHV1 (gB1) as antigen, revealed that all adult individuals possessed antibodies against gB1 (Fig 2A). Interestingly, even adult African elephants reacted positively in this assay, which can be attributed to the close antigenic relationship (89% amino acid identity) of gB1 and EEHV6-gB (gB6). In contrast, horses, which are expected to be resistant to EEHV-infections remained seronegative. Similarly, the sera of two young elephants from Germany who previously had succumbed to EHD did not reveal such antibodies. Interestingly, two of Zurich elephants at risk (Omysha; Umesh) had detectable levels of anti-gB1 antibodies in their sera, whereas the third animal (Ruwani) appeared seronegative.

**Fig 2.**
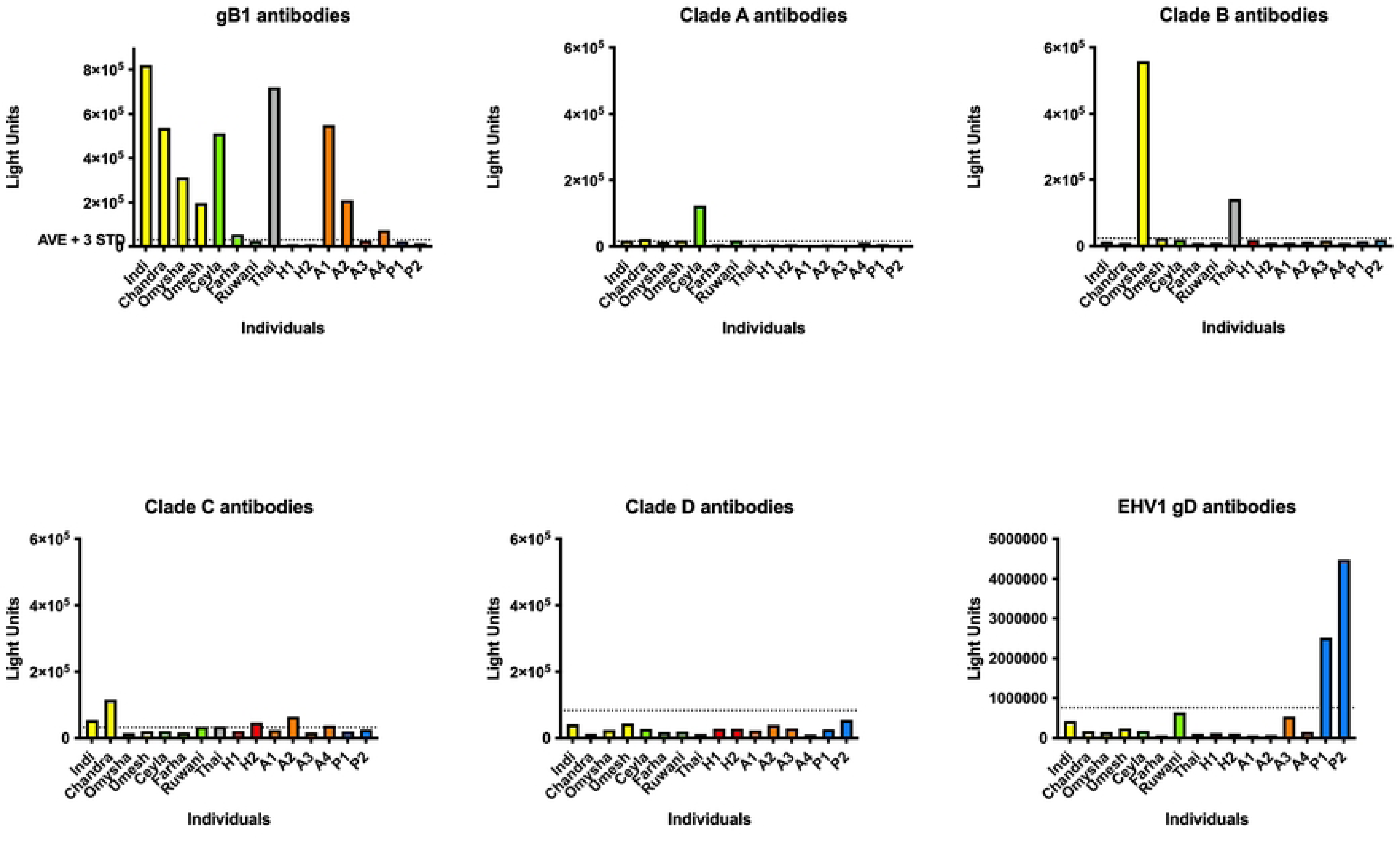
Prior antibody responses against EEHV1 antigens. LIPS assays were performed to detect antibodies against various EEHV1 antigens in the sera of elephants and horses. In each panel, the individual elephants are lined up on the x-axis, whereas the reaction of their sera is given in relative light units (RLU) on the y-axis. Yellow bars: Members of the Indi family; green bars: members of the Ceyla family; grey bar: Thai, most recent outside acquisition to the herd, introduced in 2014; red bars: sera from previous EHD cases in the Hagenbeck zoo, Hamburg, Germany; mandarin-colored bars: African elephants from the Basel zoo, Switzerland; blue bars: horse sera, which were used to determine the negative cut-off value of the test. **Panel A**. Antibody responses against the conserved glycoprotein gB1; **Panel B**: antibodies against ORF-Q clade A; **Panel C**: antibodies against ORF-Q clade B; **Panel D**: antibodies against ORF-Q clade C; **Panel E**: antibodies against ORF-Q clade D; **Panel F**: antibodies against equine alphaherpesvirus 1 gD.

Next, the elephant sera were analyzed against ORF-Q clade A, B, C, and D antigens (Fig 2B- E). In general, the antibody levels against the ORF-Q antigens were considerably lower than against the conserved glycoprotein B. However, at least one individual (Indi) reacted clearly above background levels against ORF-Q clade A antigen, whereas two individuals (Omysha and Thai) were clearly seropositive against ORF-Q clade B antigen. Moreover, at least two of the individuals at risk (Indi, Chandra) reacted positively against ORF-Q clade C antigen. Interestingly, two of our control sera (one of the German individuals who had died from EHD and one African elephant) also reacted above the background level. Although at least two individuals (Indi, Ceyla) had been present during a previous outbreak of an ORF-Q clade D virus, none of the individuals at risk was presently deemed seropositive against ORF-Q clade D antigen.

As all negative cut-off values were calculated by using horse sera, it seemed sensible to analyze the same panel of sera against an antigen (glycoprotein D, gD), which is suitable to detect antibodies against equine alphaherpesvirus 1 (EHV-1), a highly prevalent pathogen of horses. Indeed, both horse sera tested reacted positively, whereas all elephant sera had no detectable antibodies against the equine viral antigen (Fig 2F).

Together, these serological analyses confirmed both the presence and replication of various EEHV1 subtypes among the animals at risk.

### Viremia upon EHD

On Thursday, June 16, 2022, all three of the monitored animals tested negative for both EEHV1 and EEHV4, but one week later, on June 23, Umesh tested EEHV1-positive with a ΔCt value of +3.8. Samples from the two following days confirmed its increasingly viremic status with ΔCt values of -0.75 and -2.75, respectively. Eventually, the value of the same animal dropped to -4.1 and to -4.65 before coming back to +0.5 at the day of its death due to EHD, five days after initial detection of viremia (Fig 3). The other two animals remained EEHV1-negative until slight viremia was first detected on June 26. Both went back to negative in their next two samples. On July 3, Omysha’s ΔCt value was +6.15 and, after a brief recovery dropped to -1.4 before the animal died 10 days after initial detection of viremia. Beginning from June 30, Ruwani went forth and back with short periods of viremia. Eventually, the ΔCt also dropped to negative values and this animal was also lost to EHD within four weeks (Fig 3).

**Fig 3.**
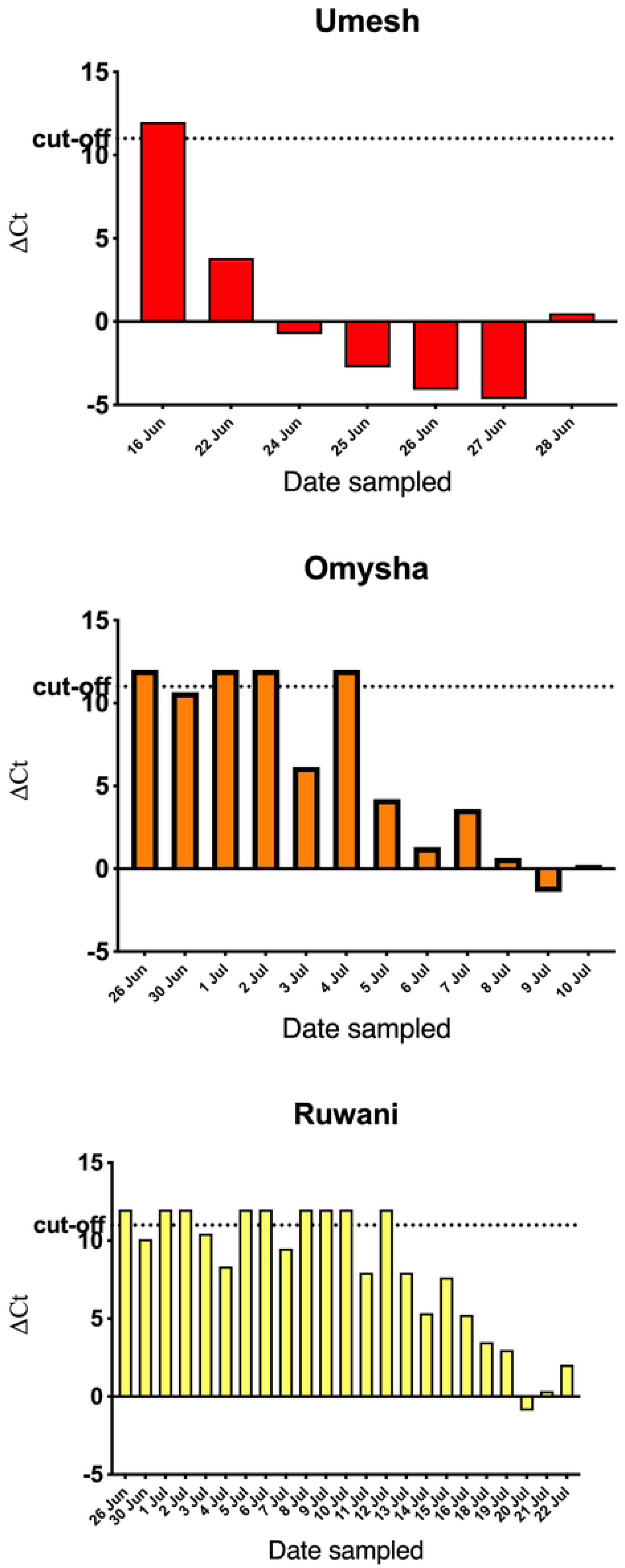
Relative EEHV1 loads in three animals before and throughout development of EHD. Delta Ct values are shown on the y-axis, whereas the date of blood sampling is given on the x-axis.

### Analysis of the causative EEHV

Sequencing of the E36/U79 locus

The above used TaqMan real-time PCR for detection of EEHV1 DNA targets the U41 gene (encoding the major DNA-binding protein; MDBP) and does not discriminate between the EEHV1 subtypes A or B. Being aware of previous circulation of both subtypes among the elephants at Zurich zoo (24), a potential antiserum treatment was considered. Therefore, it was of immediate relevance to identify the subtype of the present virus. A conventional PCR targeting the E36/U79 locus was therefore initiated and the resulting PCR product was sequenced. As shown in Fig 4, the resulting sequence obtained from an early sample of animal Umesh was 100% identical to Xian’s previous EEHV1A isolate, whereas it showed (in between the primers) 11 different nucleotides in comparison to Omysha’s previous EEHV1B isolate.

**Fig 4.**
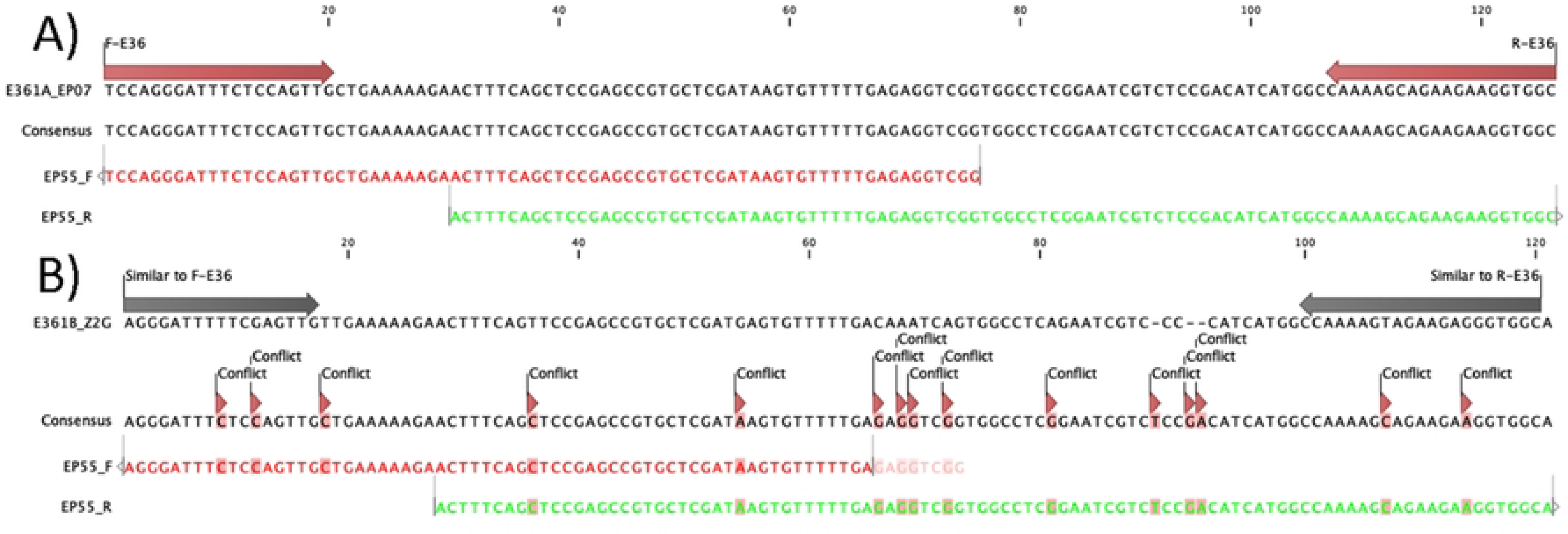
Sequence alignments. Alignment of the present E36/U79 sequence (Umesh isolate, EP55) against previous ones. (A) Comparison against the 1A subtype (EP07 isolate). (B) Comparison against the 1B subtype (trunk wash from Omysha).

### Analysis of the E54/EE51 vOX2-1 locus

As the vOX2-1 locus has been reported to be highly variable among strains of EEHV1 (up to 15%) but remains identical in epidemiologically linked cases of EHD (33), we amplified an 872 bp DNA fragment from a blood sample of Omysha by conventional PCR. Sanger sequencing of the PCR product yielded 840 nucleotides, which were compared against the previously determined vOX2-1 sequence obtained from EP07, who died from EHD in the Zurich zoo in 1999 as well as against an American (Kimba, 2004) and a European EEHV1A strain (Raman, 2009) and rooted against the subtype 1B prototype (Emelia,2006)(10, 11). On the nucleotide level, the vOX2-1 sequences of the present Umesh isolate (2022) shared only 92.34 – 93.84% identity with any of the compared sequences, corresponding to a minimum of 54 to a maximum of 67 nucleotides per sequence. The closest relationship was observed with the European 1A prototype (Raman), the least with the American 1A prototype (Kimba) (Fig 5). The present isolate (EP55) branched with an unrelated European isolate, thus clearly differing from the previous EHD cases in the zoo.

**Fig 5.**
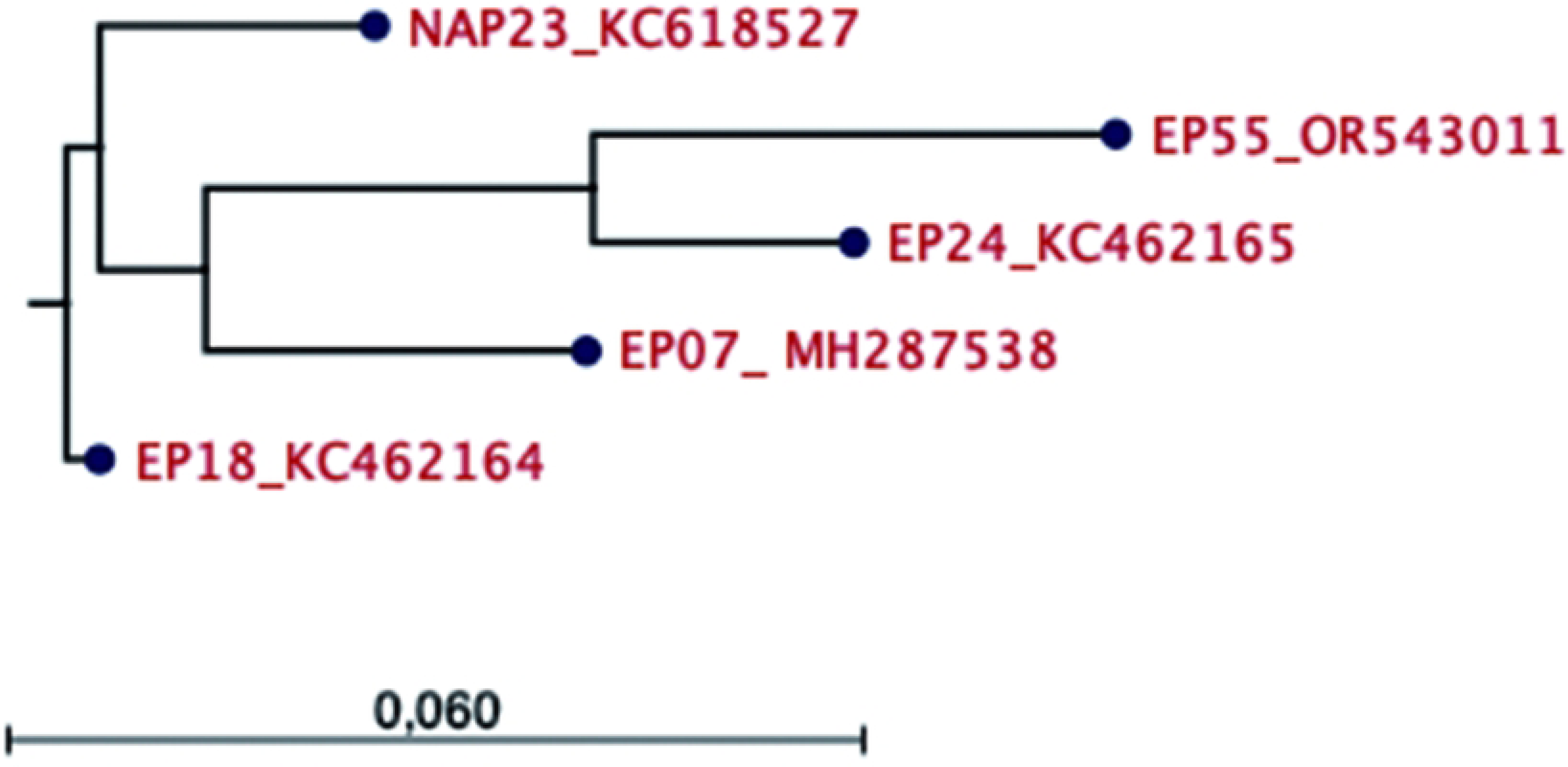
vOX2-1 phylogenetic tree. A phylogenetic tree was obtained from the alignment of the nucleotide sequences of the vOX2-1 locus (E54/EE51) from the American prototype of EEHV1A (NAP23), the European EEHV1A prototype (EP24), the European prototype of EEHV1B (EP18) in comparison to the corresponding sequences of the present case (Umesh, EP55) and a previous case of EHD in the Zurich zoo (Xian, 1999, EP07). Each strain is identified by its case number followed by its GenBank accession number.

### Determination of the full viral sequences from all three cases

Eventually, the full sequence of the present EEHV1A virus was determined from material obtained from Umesh. All raw sequencing data generated were uploaded to the Sequence Read Archive (SRA) as BioProject ID: PRJNA1052582. The sequencing data obtained from samples from Omysha and Ruwani was less complete but by comparing the contigs obtained with the Umesh genome, 27’488 and 4679 NNNs were assembled in the two incomplete EEHV genomes, respectively. Excluding non-sequenced fragments, the genomes had a high similarity, e.g., only 56 nt mismatch between Umesh and Ruwani. Therefore, the main focus of our further analysis was on Umesh’s complete genome.

### Overall size and genome structure of EEHV1A (Umesh, EP55)

The size of the assembled single complete intact genomic scaffold of Umesh (EP55) obtained by a virion enriched next-generation Illumina paired-end procedure was 179,847-bp (GenBank accession number OR543011). This included two near identical copies of the 2.9- kb direct terminal redundancy (TR), one located at each end. Most other EEHVs that we have been involved with lacked a second copy of the TR most likely because they were carried out without the virion enrichment step. Probably the most distinctive features of EEHV1 genomes are the putative origin of lytic DNA replication (ori-Lyt, 1190-bp), the two most likely nuclear immediate-early transcriptional control proteins E40 (ORF-K, 743-aa) and E44 (ORF- L, 1209-aa) and the largest non-core likely tegument protein E34 (ORF-C, 1886-aa), which are all present and intact with close to the same inverted repeat stem structures and amino acid sequences as those found in all previously studied EEHV1 genomes. Other unique features of EEHV1, include the E4 (vGCNT1), E47 (vFUT9), U73 (OBP), three versions of vOX2-like proteins E24 (vOX2-3), E25 (vOX2-2) and E54 (vOX2-1), as well as U27.5 (RRB) and U48.5 (TK) plus a total of 10x vGPCR-like proteins E3 (vGPCR6), E5 (vGPCR5), E15 (vGPCR4), E20 (vGPCR4A), E21 (vGPCR4B), E26 (vGPCR3), U12 (vGPCR2), U51 (vGPCR1), E50.5 (vGPCR9) and E59 (vGPCR10) and a large family of 18 RAIP-like 7xTM domain receptors are also all present. With just a small number of exceptions, the same applies to the majority of the rest of the expected total of about 120 genes of the virus. Previous assessments of predicted ORFs and of in-frame splicing plus exon donor and acceptor motifs and intron sizes and boundary patterns for the rest of the protein coding genes were originally based largely on other known mammalian herpesviruses and especially on consistency across multiple independent EEHV1 strains.

Almost 40 of the genes encoded by EEHV1 that are also found (albeit as highly diverged versions) within the majority of other mammalian herpesvirus sub-families are referred to as “core” genes with an applied U number nomenclature system derived from that used originally for human HHV6 genome analysis. In comparison, all of the remaining 78 protein encoding genes that are novel and not found in any other herpesviruses are given a sequential unique E number nomenclature system based on that originally applied to the prototype EEHV1 (Kimba) genome (11). For individual protein names, where feasible we have tried to apply the most universally agreed upon three-letter abbreviation for those proteins when some previously available functional or homology data is available from other well-studied herpesvirus species, and have otherwise used a combination of either an extended ORF-A to ORF-S system as originally suggested by Ehlers (17) or just left them with the same E number as for the designated gene name where little else is known for novel *Proboscivirus*- specific proteins.

### Comparison with other EEHV1A and EEHV1B genomes

Previous experience in EEHV1 whole genome analysis has revealed that all individual independent strains display unexpectedly high levels of overall genetic divergence associated with a large subset of hypervariable genes. Across the entire population of EEHV1 genomes the most striking feature is that these numerous significantly diverged variable genes nearly always cluster into cladal subsets that can be designated as specific subtypes. It is also evident that most (but not all) EEHV1 variable genes often show these large and complex differences at both the protein (amino acid) level as well as at the DNA (nucleotide) level. Fig 6 shows the results of a search for intergenomic similarities which revealed that the closest matched of the other nine complete EEHV1A genomes contained in the GenBank database is the Indian strain EEHV1A (IP43) [MN366291]. The latter differs from Umesh (EP55) by 2.7% at the nucleotide level (4,856-bp mismatch) as determined by direct pairwise comparisons using the VIRIDIC tool (34). It also differs from our prototype EEHV1A (Kimba) genome [KC618527] by 4.6% (9,712-bp mismatch) and is least closely related to the EEHV1A (IP143) genome [MN366294] with 7.5% divergence (13,488-bp mismatch). All of the other EEHV1A intact genomes fall within the range between IP43 and IP143. For the single prototype chimeric sub- species EEHV1B (Emelia) genome [KC462165] from *Elephas maximus* the divergence is 8.2% (14,747-bp mismatch) and for the single orthologous prototype EEHV6(NAP35) genome [MZ822422] from *Loxodonta africana* the divergence is 26.2% (47,120-bp mismatch).

**Fig 6.**
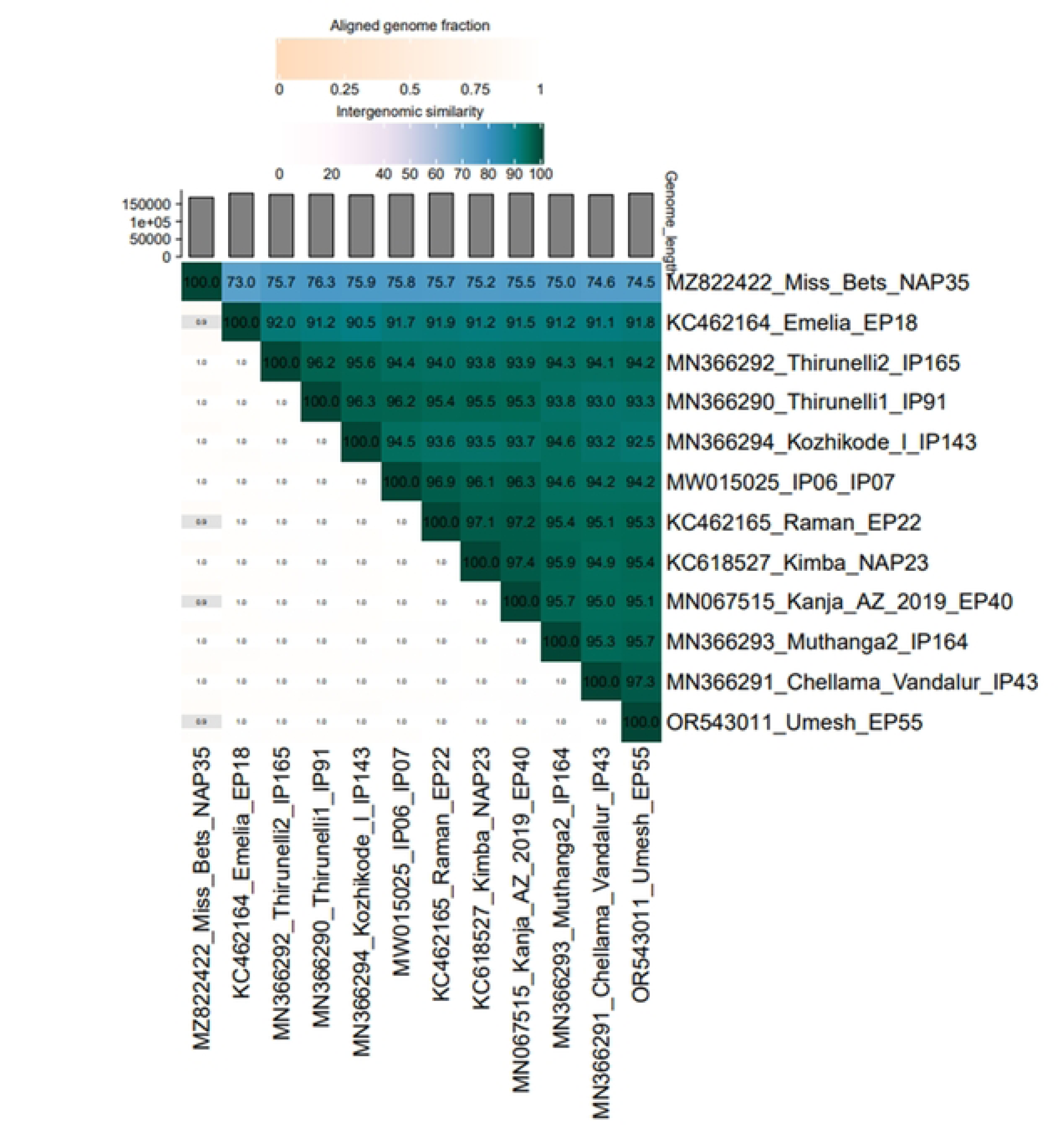
Similarity heatmap. BLASTN alignment of the Umesh (EP55) genome against closely related EEHV genomes in the GenBank database using the VIRIDIC tool. The heatmap shows the similarity values for each genome pair, the fraction of genomes aligned and the genome length.

After detailed BLASTX and BLASTP comparisons of all 122 genes individually at both the DNA and protein level relative to all other sufficiently well-characterized EEHV1 strains, the genome of Umesh proved to display similar overall features and gene organization as well as the typical sporadic non-uniform patterns of divergence. It is noticeable that Umesh does not actually contain any extra or novel genes not seen before in at least a subset of EEHV1 strains. But despite that statement, this does not imply that it contains exactly the same set of genes or gene subtypes as any or all other EEHV1 strains. In fact, there are indeed a small number of overall differences in both gene content and genome organization and no other previously evaluated strains have the exact same subtype patterns for the many designated hypervariable genes. Just like all the other strains evaluated Umesh consists of a complex mosaic pattern of mostly well-conserved genes intermingled together with a distinctive and unique pattern of subtypes for the moderate and highly variable ones.

In general, although our earlier analysis at several short PCR loci predicted that Umesh would most likely prove to be an example of an EEHV1A not EEHV1B subspecies, to be certain that this was valid across the whole genome, we needed to evaluate all 12 individual gene loci that almost universally and consistently define the relatively rare chimeric EEHV1B sub-species as distinct from the more commonly encountered non-chimeric EEHV1A sub-species. These all map within three major non-adjacent multi-gene chimeric domains in the central regions of the genome referred to as CD-I (gB/POL), CD-II (ORF-J/gN/gO/gH and TK) and CD-III (UDG, gL/ORF-O/ORF-P/ORF-Q), which together encompass nearly 13-kb and with one exception (POL) differ by at least 10% at the protein level between the prototypes of the EEHV1A (Kimba) or EEHV1A (Raman) sub-species and the EEHV1B (Emelia) sub-species (5, 10).

Although EEHV1B (Emelia) is the only completely sequenced member of the 1B sub-species, there are eight other partially characterized EEHV1B genomes, namely NAP14 (Kiba), NAP33 = NAP34, NAP38, NAP45, NAP47A, NAP49 and IP93 that almost universally also have B-specific characteristics at all of these same loci. Furthermore, there are also three other frequently analyzed gene loci that display universal B sub-species features, but only differ by less than 10% from the 1A prototype e.g. U51 (vGPCR1), U71 (MyrTeg) or U73 (OBP) that have all proven to have universally linked B-subtype versions also, plus four others that are only partially linked to the rest of the 1B diaspora e.g.E24/E25 and E30/E31. For example, only NAP14 and NAP49, but not Emelia, NAP33 or NAP45 have the highly diverged subtype-B pattern for the E24/E25 locus and only Emelia, NAP14, and NAP49, but not NAP33 or NAP45 have the B-subtype pattern for the E30/E31 locus. Finally, there are also three variable genes among the novel 7 x TM domain proteins that each split cleanly into three distinct subtype clusters, but that have no linkage or obvious connection whatever to the classic 1A versus 1B chimeric patterns nor to each other (E3/vGPCR6), E5/vGPCR5 and E26(vGPCR3).

### Comparison with main segment EEHV1 variable gene subtype assignments

Full evaluation of Umesh (EP55) shows that it does not contain even a single gene locus that exhibits a 1B-like pattern and it does also contain the single small E36A (ORF-N) vCXCL chemokine-like protein that is unique to EEHV1A strains and absent from all EEHV1B strains. In contrast, the Umesh assigned subtypes at each of the 19 variable gene loci included above are as follows: E3 (vGPCR3 = A1; E5 (vGPCR5) = C; E24 (vOX2-3 = A, E25 (vOX2- 2) = A, E26 (vGPCR3) = B; E30 = C1, E31 = C, E39 (gB)= C1, E38 (POL) = A, E35 (ORF- J) = A, U46 (gN) = C1, U47 (gO) = A1, U48 (gH) = D, U48.5 (TK) = A, U51 (vGPCR1) = E, U71 (MyrTeg) = A, U73 (OBP) = A. U81 (UDG) = A, U82 (gL) = A1, E37 (ORF-O) = C, E38 (ORF-P) = E, and E39 (ORF-Q) = C. A detailed comparison of this information for Umesh with the equivalent subtype cluster assignments for the same set of 22 variable genes from nine other selected EEHV1 strains is presented in the upper section of supplementary **Table S1**. The particular examples chosen to be shown here include the original three complete genome protoypes from GenBank of EEHV1A (Kimba), EEHV1A (Raman) and EEHV1B (Emelia), plus accumulated gene locus data from six of the best characterized but incomplete EEHV1A genomes determined previously by conventional PCR amplification- based DNA sequencing of older lethal EHD cases carried out by our Johns Hopkins University Laboratory group. Specifically, this data was derived from one sample from India (IP11), one from Europe (EP07, Xian) and four from the United States (NAP18, NAP26, NAP17 and NAP21). With the exception of the prototype EEHV1A (Kimba) strain, which (although it does have three Cs within CD-II) was otherwise designated to have just A or A1- subtype variable genes across the entire genome, all other non-1B diaspora strains fall into complex mosaic patterns like this with largely unlinked variable gene subtypes.

The most useful region to evaluate here has proven to be the combined ORF-O/ORF-P/ORF- Q complex (part of CD-III) for which we recognize 13 different triplex patterns, with the same C-E-C subtypes as in Umesh (EP55) being found also in 11 out of 52 other EEHV1A or EEHV1B cases that have been fully evaluated previously at this locus, namely Raman, EP14, IP43, NAP16, NAP24, NAP43, NAP73, NAP75, NAP88, NAP91 and NAP93. Although only two of those are available as complete genomes (Raman and IP43), none of them occur within known EEHV1B strains. In addition, only one of the seven India cases evaluated here (IP43) has that same C-E-C group pattern and only two others have been identified among the relatively few European EEHV1A cases examined in detail so far (i.e. Raman and EP14 but not EP07 or Umesh). Beyond those specific loci discussed above that are either fully or partially related to the 1A/1B sub-species chimeric domain patterns there are also another three variable genes among the novel 7 x TM domain proteins that each split cleanly into three distinct subtype clusters, but that have no linkage or obvious connection whatever to the classic 1A versus 1B chimeric patterns nor to each other i.e. E3 (vGPCR6), E5 (vGPCR5 and E26 (vGPCR3).

### Comparison with other EEHV1 R2-segment gene arrangements

Similarly, many of the 11 potential gene coding positions located within the short 10.5-kb R2- segment of all EEHV1 genomes display high levels of variability and subtyping that have proven to be either completely unrelated to the linked 1A/1B chimerism diaspora pattern explained above (or else are very highly scrambled). These latter non-1A/1B diaspora related hypervariable gene loci all map within the novel *Proboscivirus*-specific segments located near both ends of the linear EEHV1 genomes. Amazingly, although the R2-segment mapping between E47 (vFUT9) to E55 (vIgFam3) encompasses just 5% of the length of the EEHV1 genome, it has multiple levels of additional complexity that together provide a large fraction of the overall EEHV1 genomic variation. Furthermore, we have interpreted that this part of the complexity within the R2-segment involves multiple alternative two or three-gene “cassettes” that define eight distinct clustered R2 sub-groups designated A, B, C, D, E, F, G and H. These latter cluster groups represent a major determinant of EEHV1 genomic organization within non-1B strains.

The presence or absence of these eight types of inserted multigene “cassettes” as well as which of three different potential genomic locations where they are inserted at largely defines each R2 group. However, unlike the 20 known available “optional” R2 family genes present in the “cassettes”, which are each found in just a small subset of the population, there are also eight remaining “constant” R2-segment genes, that are almost always all present, but are also mostly highly diverged and form many additional subtype clusters of their own. The latter include two interesting single genes E47 (vFUT9) and E54 (vOX2-1) and six other members of R2-gene families that are largely common to all strains. Therefore, although each individual EEHV1 genome usually contains a total of just 11 genes (often including an average of two or three fragmented or inactivated ORFs) within the R2-segment, the overall known population of 37 distinct independent EEHV1A plus EEHV1B virus strains that have been evaluated fully across the R2-segment encodes as many as 28 different total alternative available genes.

These other “potential” R2-genes include seven paralogous versions of RAIP-related vGPCR- receptors E48 (vGPCR7), E56 (vGPCR8), E50.5 (vGPCR8/vGPCR9/vGPCR13), E59 (vGPCR10), E62 (vGPCR11), E56 (vGPCR12) plus 18 paralogous diverged versions of (mostly) membrane-bound immunoglobulin family (vIgFam1 to vIgFam15) proteins, of which some are detectably related to host cell protein CD48, and a third family of four other rather non-descript membrane proteins (E49, E51, E64 and E68). The presence (+) or absence (-) plus the relative location and subtype designation of twenty representative genes found within the R2-segments of both Umesh and the same set of nine other selected comparative examples of EEHV1 genomes are summarized in the lower section of supplementary **Table S1.** Note that only four of the eight known organization patterns of alternative R2-segment insertion cassettes are listed here (i.e. A, B, D and F groups, but not C, E, or G). From amongst the known total of 28 possible alternative R2-genes, Umesh (EP55) most closely matches the Ganesh (NAP26) and IP43 patterns by having a typical group-D E59/60/61 triplex cassette and encodes the following 11 proteins overall with their assigned subtypes as shown i.e. E47 (vFUT9) = B, E50.5 (vGPCR9) = C, E51(membrane) = D1 (frame-shifted), E52 (vIgFam2) = deleted, E59 (vGPCR10) = D, E60 (vIgFam6) = +, E61 (vIgFam7) = + (frame-shifted), E52.5 (vIgFam2.3) = deleted, E53 (vIgFam2.5) = frame-shifted, E54(vOX2-1) = o and E55 (vIgFam3) = A2. Unlike the other R2 genes described here neither the E60, E61 or E53 proteins show any significant subtype variability, whereas in dramatic contrast the 43 examples of E54 (vOX2-1) proteins evaluated despite collectively exhibiting up to 15% overall variability at the aa level they do not cluster into any readily definable subtype groupings. Furthermore, although there are just two tight major subtype clusters of E47 (vFUT9) in which the prototype EEHV1A (Kimba) version (subtype-A) and the prototype EEHV1B (Emelia) version (subtype-B) differ by as much as 32% at the protein level, this particular additional example of subtype clustering is apparently not related in any way (or at least not linked to) the classic 1A/1B subspecies diaspora. In fact, from among the 35 total E47 (vFUT9) enzyme-encoding genes evaluated there are more examples of the B- subtype (24x) than of the A-subtype (11x) with both seemingly randomly assorted among the otherwise nominally A sub-species versus B sub-species genomes.

Overall, we consider the eight different two or three-gene inserted “cassette” types to be the most definitive way to classify EEHV1 genome patterns within the R2-segment, and note that once again the Umesh strain clusters within the largest set (= group-D) here along with ten other group-D cases (namely Daisy, IP43, EP07, EP20 = EP21, NAP30, NAP31 and NAP41 = NAP47), including three from Europe (EP07, EP20 = EP21), but just one from India (IP43). Moreover, there are no known group-D inserted cassette patterns (i.e. E59/E60/E61) that fall amongst the six known EEHV1B genomes that have been fully evaluated across the R2- segment. However, there are several more anomalous situations where a total of 10 virus strains have either just a two-gene cassette (groups-C, -E and -G) and four others have both a B-group like three-gene cassette plus a two-gene H-group cassette. The next two most abundant “cassette” patterns beyond the D-group are 7 x As and 6 x Cs, with the remaining 15 examples being 4 x F, 2 x E, 3 x C, 2 x G and 4 x H.

Finally, in terms of detailed gene-by-gene analyses, there are two alternative published physical genetic maps of prototype EEHV1A genomes that Umesh (EP55) can be compared to i.e. those of EEHV1A (Raman) from Wilkie et al (10) and EEHV1A (Kimba) from Ling et al (11), but note that they are drawn in opposite orientations and use different gene nomenclature systems. Furthermore, the Umesh genome described here and all other intact annotated EEHV1, EEHV2A, EEHV5B and EEHV6 genomes that we have submitted to GenBank since the original EEHV1A prototype have the same orientation and nomenclature as used for Kimba (11), except that just the R2-segment itself had to be switched to the extreme left-hand-side once the actual linearizing packaging cleavage site was determined (10). Curiously, none of the GC-rich branch *Probosciviruses* EEHV3A, EEHV3B nor EEHV4A or EEHV4B complete genomes even have the R2-segment (18).

Overall, the sequencing results confirmed that all three elephants had been hit by the same novel strain of EEHV subtype 1A. More specific Umesh (EP55) data relating to both E39 (ORF-Q) serology and three viral-encoded enzymes of interest in the context of epidemiology and antiviral treatment are presented below.

### Analysis of the E39/EE3 ORF-Q locus

Antibodies against a particular ORF-Q clade have previously been associated with a mild course of EEHV1 infection, whereas their absence have been linked to severe EHD (7). Having established that the present virus was different from the two previously observed ones, it was of interest to determine its ORF-Q (E39/EE3) sequence. Therefore, the E39 locus of the virus in Umesh’s samples was sequenced and the translated amino acid sequences were aligned to generate a phylogenetic tree (Fig 7). According to this analysis, the Umesh (2022) isolate was nearly identical with other ORF-Q Clade C isolates (EP14 and 24; NAP43, 73, 75; IP43) but clearly different from Clades A (NAP11, 16,18, 23, 26, 32, 39, 41, 47, 80; IP11, 164), D (EP07 (Xian), EP21; NAP20,29,30,31,72, IP07), B (NAP45, 49), E (NAP21) and F (NAP17) and also from EEHV6 (NAP35), whose ORF-Q sequences were used for routing of the tree. Unfortunately, these results came too late to consider a serum transfer from one of the Clade C-positive animals. Together, these epidemiologically relevant data stated that a previously unrecognized, newly detected strain of EEHV1A had been the cause of the present EHD outbreak.

**Fig 7.**
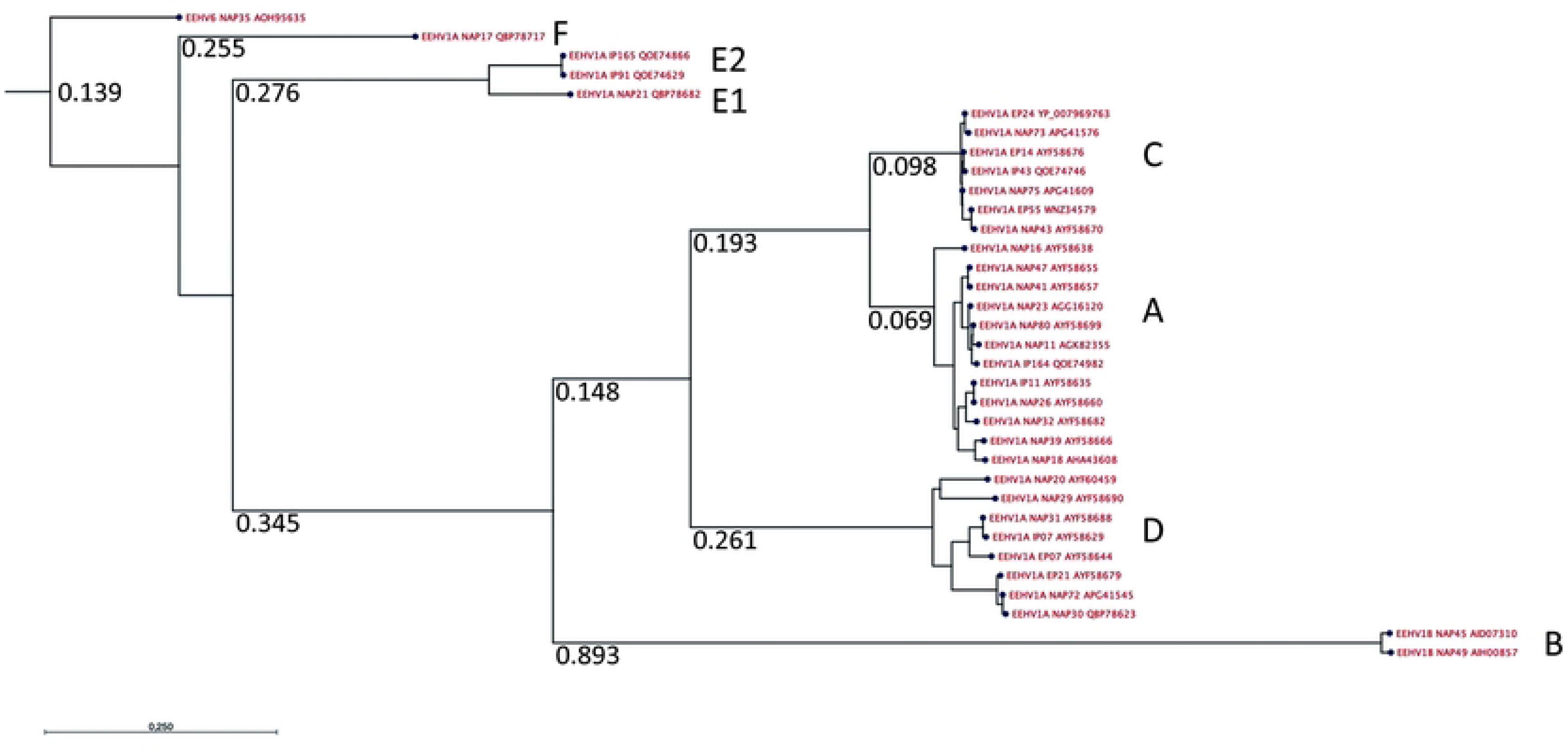
ORF-Q phylogenetic tree. A maximum likelihood phylogenetic tree was obtained from the alignment of ORF-Q amino acid sequences using the Neighbor Joining method and 100 replicates to perform bootstrap analysis. The branch length is indicated for major branches, dividing the tree into the known clades A through F. Each strain is identified by its case number followed by its GenBank accession number.

### Analysis of the U48.5/EE7 ETk locus

As the ETk may contribute to the success of antiviral treatment with nucleoside analogues, it was of interest to compare the predicted amino acid sequence of the present EEHV1A isolate against previously known ETks. Indeed, the aa sequence was highly conserved among reference EEHV1A isolates, displaying just one aa difference (H278N) against the sequence from the European EEHV1A prototype (Raman) and two aa (V152I and H278N) against the American prototype (Kimba).

This strong aa conservation of the predicted enzyme among only remotely related EEHV1A strains strongly speaks for a biological role, although its *in vitro* and *in vivo* activity against the presently used antiviral compounds remains to be determined.

### Analysis of the U69 ECPK locus

Similarly, the present ECPK aa sequences were compared against those of other known EEHV isolates. Not unexpectedly, those aa sequences were highly conserved among EEHV1A strains. Not a single aa difference from the present sequence was found in the sequences of the European EEHV1A isolates Raman or Plai Kiri. One single aa exchange (T409A) was detected upon comparison against the EEHV1B isolate Emelia. The ECPK sequence Kimba comprised two aa differences, namely T408P and T409P. Since all these differences map outside of the predicted catalytic domain of the enzyme (aa 166 to 306), it will be of interest to address the biological functionality of it in the future.

### Analysis of the U38 DNA polymerase locus

The viral DNA polymerase enzyme, encoded by U38, is considered essential for viral DNA replication. Phosphorylated nucleoside analogues may either block the enzyme to terminating its activity or integrate chain-terminating molecules into the growing DNA, which blocks further elongation. It was, therefore, of interest to compare this locus in between our three isolates as well as against other published amino acid sequences in the databases. Notably, the sequence conservation among our three isolates at this locus was 100%, whereas two to eight amino acid changes were observed upon comparison of our translated sequences against the amino acid sequences from 10 previously published cases. Interestingly, a single one of these differences in our isolates opposed to high conservation among all others, i.e., Leu 792 in our sequence compared to Isoleucine at this position in all ten previously published isolates (L792I). Moreover, this seeming mutation mapped to the center of a conserved domain, which had been associated with predicted enzymatic activities, including elongation, DNA-binding and dNTP binding. All other differences had been previously described for other U38 proteins.

## Discussion

With two prior cases of EHD in the zoo, the prevalence of EEHV1A, EEHV1B, and EEHV4 among the adult animals, and the presence of two seronegative animals in the susceptible age, the risk for another outbreak of EHD was considered high.

During four years of weekly screening, seven instances of low grade, transient viremia were detected, each single one caused by EEHV1, while viremia due to EEHV4 was never detected. These findings confirmed earlier observations of the presence and active circulation of EEHV1 among these animals.

EEHV serology has only recently established its proper role in addressing EEHV prevalence among elephants (2, 7, 14). Particularly LIPS assays have allowed to discriminate between antibodies that cross-react among various EEHV types or strains and other antibodies that represent a specific immune response against a particular virus strain (7). We selected LIPS assays using the conserved, cross-reactive glycoprotein B- (gB) and the strain-specific ORF-Q-antigens, respectively, to assess the serological status of the animals against EEHV. The use of ProteinA/ProteinG-beads in the LIPS assays brings a great advantage for the veterinary medical purpose with it because different animal species can be addressed using the same assay. We took this advantage in our case by using horse sera to determine the negative cut- off values for all of our LIPS assays.

The sera of all adult elephants had detectable levels of antibodies against glycoprotein B of EEHV1A (gB1A). Moreover, four sera from African elephants, kindly provided by the Basel zoo in Switzerland, yielded also varying degrees of antibodies against gB1A. The published amino acid sequence of the EEHV6 glycoprotein B (gB6) shares 88 to 89% amino acid identity with gB1 of the subtype 1A strains from Kimba and Raman, respectively.

Interestingly the amino acid sequence of subtype 1B (Emelia) shares only about 85% identity with either EEHV6 (NAP35) or EEHV1A (Kimba, Raman). Consequently, we assume that the gB-antibodies detected in the sera from the African elephants were cross-reacting and due to subclinical infection with EEHV6.

The following observations were relevant for the three young animals at risk of EHD: At eight years of age and with a history of shedding EEHV1B as well as several occasions of transient viremia, the seropositive status of Omysha could be attributed to having mounted its own active immune response against EEHV1. At two years of age and without previously detected episodes of viremia, the seropositive status of Umesh was somewhat unexpected. Maternal antibodies were expected to be quite low at this age but as half-brother and playing companion to Omysha, it is possible that this individual had already mounted its own immune response against EEHV1 under the previous cover of maternal antibodies. The seronegative status of Ruwani at the age of five years was a matter of concern. Neither traces of maternal antibodies could be detected any more nor an active immunity seemed to have developed, although three episodes of viremia had previously been recorded.

LIPS assays using ORF-Q antigens revealed much fewer seropositive animals than gB- serology. As the subtype EEHV1B had been observed to be shed from the trunk of Omysha (24), it was not surprising that this animal showed the highest reaction among the members of the herd against the ORF-Q-B antigen. In contrast, the other young animals at risk did not have such antibodies nor were they seropositive against ORF-Q antigens A, C, or D. Of the older members of the herd, at least one elephant cow (Ceyla) had significant antibody levels against ORF-Q-A, whereas two elephant cows (Indi; Chandra) had preexisting antibodies against ORF-Q-C. Interestingly, none of the animals seemed to have significant levels of antibodies against ORF-Q-D antigen, although the isolate from a previous case of EHD in the zoo (EP07, 1999) had been classified to belong to ORF-Q Clade D. These results suggested that antibodies against ORF-Q antigens may be short-lived and that the young animals were indeed at risk upon an EEHV outbreak.

Thanks to the preventive viremia-screening program, the 2022 outbreak was detected early. Despite of various treatment attempts (to be reported elsewhere), the virus burden in the bloodstream increased dramatically and 6 days after viremia was first detected, Umesh succumbed to EHD. The second animal (Omysha) rebounded twice from viremic to aviremic and back but then still succumbed to EHD. The third animal (Ruwani) had its first episode of viremia at the same day as the second animal (Omysha) but had a survival time of three weeks after the first episode of viremia. Only upon its fifth episode of viremia, the virus burden in the blood stream increased with almost each consecutive sample and the animal eventually succumbed to the disease.

Once the outbreak was ongoing, it was of immediate interest to subtype the causative virus more closely. Indeed, sequencing of the E36/U79 locus ruled out any involvement of a subtype 1B strain and identified, instead, a subtype 1A virus as the present disease-causing agent. This conclusion raised new concern about the eight-year-old Omysha, which had been found seropositive for antibodies against the ORF-Q-B antigen but not against other ORF-Q antigens. Moreover, the possibility to use her antibody-rich serum for a potential treatment of the other animals at risk was now ruled out.

Subsequently, it was inviting to speculate that the previously detected 1A strain, dating back to 1999 and 2003 had re-emerged after long years of latency, which had been the case in Berlin, with Shaina Pali and Ko-Raya (EP23 and EP25) being identical to Plai Kiri (EP14) from 11 years earlier (1). Yet, a comparison of E54/EE51 sequences, including the nucleotide sequences of the 1999 (EP07) isolate and the present virus revealed significant differences. Thus, the present EEHV1A virus was newly detected throughout this outbreak, raising the question about its source. These analyses during the ongoing outbreak had apparently direct effects on our risk assessment as well as treatment options.

Only once the outbreak of EHD was over, the entire EEHV1 genomes from all three cases could be determined, upon which the previous PCR analyses were confirmed. Moreover, the sequences of other genomic loci of interest became available for interpreting the course of the disease, in particular the epidemiological chain, and the failure of treatment.

Concerning the epidemiological chain, it is important to state that all three isolates had the same genomic sequence, belonging to the subtype 1A. The E54/EE51 vOX2-1 locus was 100% conserved among the three isolates, which confirmed the common origin, but was significantly different from the 1999 EP07 isolate. Analysis of the E39/EE3 locus of Umesh (EP55) identified a virus belonging to the ORF-Q-C clade. This finding was very surprising, since no such virus had previously been identified among these animals The previous EEHV1A, responsible for the death of EP07, belonged to the ORF-Q-D clade, while serological evidence suggests that the EEHV1B isolate detected in trunk-wash samples from Omysha belonged to ORF-Q-B.

The ORF-P and ORF-Q genes of EEHV1, which arose by duplication and have some residual homology, are replaced in EEHV3/4 by a single similar gene at the same location and with the same two exons (E39A, ORF-R), but with minimal or no homology to ORF-Q itself (5). Similarly, EEHV2 and EEHV5 also lack ORF-Q but do have ORF-P. Accordingly, our serology was unable to evaluate the serological status of the animals towards EEHV3/4, although EEHV4 had been detected in the trunk wash sample from Thai. Unfortunately, the ORF-Q sequence from Aishu, who died from EHD in 2003, was not available. However, the few known parts from the Aishu-virus genome, for example the U48.5/EE7 ETk locus, differ considerably from the present sequences. Overall, these sequences confirmed the presence of a new virus among these elephants.

Three different EEHV1 proteins may be involved in the susceptibility or resistance of nucleoside-based antiviral treatment against EEHV: the viral thymidine kinase (U48.5/EE7, ETk), the conserved protein kinase (U69, ECPK), and the catalytic unit of the viral DNA polymerase (U38, DNA pol).

Identities and differences, respectively, in the coding sequences within these three genes of our three isolates might not only convey about potential drug-resistance but also about the origin and sequence of transmission. If all sequences from our three isolates were identical, the source of infection must be the same for all three animals or the virus must have been transmitted from one case to the other without genomic alteration. Therefore, it is safe to assume that all three animals were infected from the same original shedder.

As its name suggests, the ECPK protein was even more conserved than ETk. Moreover, this enzyme showed *in vitro* activity, converting GCV and PCV in a dose-dependent manner, thus, making it a prime candidate for possible emergence of a drug-resistant mutant virus (1). Although a few variations against previous sequences were detected, all the differences had been identified before and mapped outside of the predicted catalytic domain of the enzyme (aa 166 to 306). Despite of the long survival time of the third case, none of the observed mutations could explain the development of a drug-resistant phenotype and ECPK. Although the details of the treatment attempts during the outbreak were not disclosed to the authors of this communication, it is possible that the U38 DNA polymerase may be naturally incompetent to incorporate certain phosphorylated nucleoside analogues into the growing viral DNA. Due to the lack of suitable cell cultures, the role of the U38 protein in EEHV- replication and its susceptibility or resistance against nucleoside analogue treatment has not yet been addressed. Still, an interesting variation was observed within the U38 DNA pol sequence, where a single mutation (L729I) in all our isolates opposed to the high conservation among ten randomly selected previous isolates. This seeming mutation mapped to the center of a conserved domain, which had been associated with predicted enzymatic activities, including elongation, DNA-binding, and dNTP binding.

Having established that all three animals had been infected most likely from the same source with the same but previously unrecognized virus, its most probable origin can be attributed to an elephant who was born outside of the Zurich zoo. Indeed, ORF-Q serology identified two individuals, Indi and her daughter Chandra, which had been ORF-Q-C-seropositive prior to the outbreak, while all other members of the herd, in particular the newest member Thai and the oldest external animal, Ceyla (born 1975 in Sri Lanka, transferred to the Zurich zoo in 1976), had been seronegative. As one of the ORF-Q-C-positive Chandra had been born and raised in the zoo, whereas her equally seropositive mother (Indi) had been born 1986 in Myanmar, was imported to Switzerland in 1988, before being transferred to the Zurich zoo in 1999), it is tempting to speculate that Indi has acquired this “novel” virus in her youth but kept it latent or did not spread it until prior to the present outbreak.

## Conclusions

Despite considerable preventive efforts and early detection of EEHV1A viremia, the Zurich zoo lost three young elephants to EHD. The reasons for the failure of the preventive measures may be summarized in three points: (1) Presence of a previously undetected EEHV1A strain among the resident elephants. Probably, the new virus re-emerged after almost 40 years of latency from one of the oldest elephants in the zoo. (2) Insufficient or absent strain-specific immunity in the affected animals. (3) Lack of efficient antiviral drugs and protective vaccines, due to the unavailability of appropriate cell cultures and animal models.

## Acknowledgments

We are grateful to the zoos in Zurich, Basel, and Hamburg for their contributions. Equally, Dr. Monika Hilbe and Prof. Dr. Franco Guscetti, Institute of Veterinary Pathology, University of Zurich are greatly acknowledged for providing necropsy tissue. We also thank Prof. Dr. Cornel Fraefel for his constant support of the viremia screening and antibody testing. Finally, we thank Andreina Schramm for sharing her EHV1 positive horse reference sera.

## Supporting information

**S1 Table. EEHV1A (Umesh, EP55) Variable Gene Subtype Comparisons**.

+ = Inserted alternative gene casettes that do not display subtyping.

- = Absence of any inserted gene casettes at these loci.

Del, fr or fs = deleted, fragmented or frame-shifted protein versions.

O = Intact E54 proteins which display up to 15% variability at the aa level, but without any clustal subtyping patterns.

* E37(ORF-O), E38 (ORF-P) and E39 (ORF-Q) proteins are also part of the CD-III chimeric domain.

\# Just four of the eight known alternative inserted three-gene cassette groups (A, B, D and F) in the R2-segment were selected for inclusion here (from a total of 30 EEHV1 strains examined).

## Notes

### Competing Interest Statement

The authors have declared no competing interest.

